# HDfleX: Software for flexible high structural resolution of hydrogen/deuterium-exchange mass spectrometry data

**DOI:** 10.1101/2021.12.09.471740

**Authors:** Neeleema Seetaloo, Monika Kish, Jonathan J. Phillips

**Affiliations:** Living Systems Institute, Department of Biosciences, University of Exeter, Stocker Road, Exeter, UK; Alan Turing Institute, British Library, London, UK

## Abstract

Hydrogen/deuterium-exchange mass spectrometry (HDX-MS) experiments on protein structures can be performed at three levels: (1) by enzymatically digesting labelled proteins and analyzing the peptides (bottom-up), (2) by further fragmenting peptides following digestion (middle-down), and (3) by fragmenting the intact labelled protein (top-down), using soft gas-phase fragmentation methods, such as electron transfer dissociation (ETD). However, to the best of our knowledge, the software packages currently available for the analysis of HDX-MS data do not enable the peptide- and ETD-levels to be combined – they can only be analyzed separately. Thus, we developed `HDfleX` – a standalone application for the analysis of flexible high structural resolution of HDX-MS data, which allows data at any level of structural resolution (intact protein, peptide, fragment) to be merged. `HDfleX` features rapid experimental data fitting, robust statistical significance analyses and optional methods for theoretical intrinsic calculations and a novel empirical correction for comparison between solution conditions.

---

Hydrogen/deuterium-exchange mass spectrometry (HDX-MS) is a widely-used analytical technique that enables the probing of protein conformational dynamics and interactions on a wide timescale, from milliseconds to hours, by monitoring the isotopic exchange at the backbone amide hydrogens with the surrounding solvent^1,2^. HDX-MS is growing in popularity as an orthogonal structural biology tool since it can analyze proteins that are not amenable to other techniques, such as antibodies^3,4^ and intrinsically disordered proteins^5,6^. In order to localize the exchange kinetics to specific regions of a protein, it is crucial to obtain sub-molecular hydrogen/deuterium-exchange data. This is currently a major limitation in the mass spectrometry detection of hydrogen/deuterium-exchange and, until recently, the structural resolution of any bottom-up HDX-MS experiment relied on the number and position of proteolytic peptides identified^5,7^. By spectral assignment of the digested peptides onto the protein sequence, the changes in deuterium uptake of each peptide can be mapped back to the primary sequence, thus giving insights into the behavior of localized regions of the protein.

Much recent interest has been directed to achieve higher structural resolution by fragmenting the digested peptides further using gas-phase soft fragmentation techniques – termed middle-down HDX-MS^8,9^. However, it is important that the fragmentation does not induce any hydrogen/deuterium scrambling (H/D scrambling)^10,11^. Methods shown to be amenable to this include electron-based dissociation methods (ExD) with HDX-MS, such as electron-transfer dissociation (ETD)^3,12,13^, electron-capture dissociation (ECD)^11,14^, and ultra-violet photodissociation (UVPD)^15,16^. These forms of fragmentation can be tuned to have analyte energy profiles that do not promote proton mobility and so result in low H/D scrambling under the right conditions^17^. This has made HDX-MS possible in topdown (fragmentation of entire proteins into smaller fragments)^13^ and middle-down (fragmentation of digested peptides into even smaller fragments)^8^ fashions, which each provide varying levels of structural resolution.

Nevertheless, presently, each of these experiments and the resulting global and local HDX-MS data can only be analyzed and interpreted separately since there is currently no widely available process that allows the combination of data at all levels^18–21^. This is especially confounding, as soft-fragmentation reactions are typically inefficient and yield sparse data which might usefully be complementary to conventional bottom-up data. In an effort to remedy this, we hereby present `HDfleX`: Software for flexible structural resolution of HDX-MS data, which allows combination of data from bottom-up, middle-down, and top-down experiments. `HDfleX` can (1) Perform nonlinear regression fitting of deuterium uptake data against labelling time for each peptide or fragment, (2) Correct for back-exchange using internal or external measures of maximal deuteration, (3) Normalize between different pH and salt conditions, (4) Calculate intrinsic rate and protection factor (*P_f_*), (5) Combine peptide and ExD fragments, and visualize the absolute deuterium uptake as histograms for individual protein states, (6) Perform hybrid significance testing using various methods of global significance testing and doing a Welch’s t-test, and (7) Visualize significant differences on difference and volcano plots.

## METHODS

`HDfleX` was written in MATLAB release 2021b (MathWorks, USA) and made into a graphical user interface with MATLAB App Designer. The software performs a series of steps summarized in Figure 1 and detailed below. An in-depth flowchart of the process can also be found in SI Figures 1-3.

**Figure 1:**
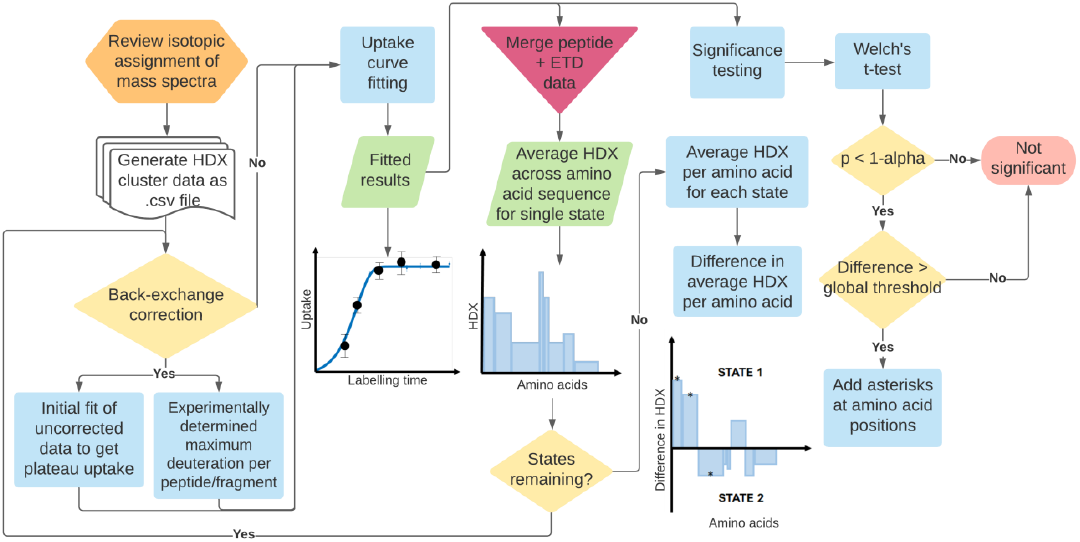
Flowchart summarizing the processes performed by `HDfleX`.

### Data import and preparation

Prior to `HDfleX` analysis, an input file containing all the information about the HDX-MS experiment needs to be created. Following review of isotopic assignments, a comma-separated variable (.csv) file is generated, which is the first user-input to `HDfleX`. The file must be comma-separated and have the nine columns in the order: protein name, sequence start number, sequence end number, sequence, modification, fragment, maximum possible uptake, mass of monoisotopic species, state name, exposure time, file name, charge, retention time, intensity, centroid. After `HDfleX` parses the input file, the user selects the protein and states to plot, type of back-exchange correction to perform (if at all), number of replicates collected, number of phases into which to fit the uptake curves, pH of reference state, pH of each experimental state, time format between milliseconds (ms), seconds (s) and minutes (min), and the time window over which the fitted uptake curve will be evaluated. In cases where the replicates are not named trivially (e.g., file_01, file_02, etc.), the user can manually assign files to the appropriate states and replicates. Charge deconvolution and deuterium uptake per peptide and ETD fragment are determined as described in Wales et al^22^.

### Uptake curve fitting and area under the uptake curve

Fitting the HDX-MS data from equilibrium experiments can provide crucial information about the exchange kinetics. We fit the data points for each peptide and ETD fragment either by interpolation or fitting to a stretched exponential. The deuterium uptake at labelling time t, is fitted to a stretched exponential given by Equation 1^23^, where *nExp* is a user-defined number of exponential phases, *N* is the maximum number of labile hydrogens, *k_obs_* is the observed exchange rate constant and *β* is a stretching factor.

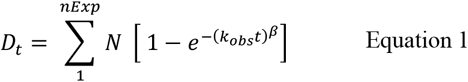

The non-linear fitting is performed using the fit function from the Curve Fitting Toolbox in MATLAB. If *nExp* = 1, *N* is fixed at the maximum number of labile hydrogens for that particular peptide/fragment if back-exchange correction has been applied. Otherwise, only an upper bound of *N* is applied. *k_obs_* and *β* are both limited between 0 and 1. The resulting fitted curve equation is subsequently evaluated and integrated over the user-defined time window to give the area under the uptake curve, HDX_Area_. In the case of data that reach a plateau (i.e., maximal deuteration achieved at later timepoints), a further variable called X_plat_ (SI Figure 4) is calculated, which is used as the upper limit of the time window of integration. This prevents significant differences from being minimized when overly wide time windows have been used for any individual peptide.

There is also an option to fit the data points using an interpolation method. Here, we used the Piecewise cubic Hermite interpolation (`PCHIP` from MATLAB’s `Curve Fitting Toolbox`). However, this method does not provide a *k_obs_* and *β*, and thus cannot be used for empirical adjustments, observed exchange rate constant and protection factor analyses.

The entire `HDfleX` workflow following uptake curve fitting, such as data flattening and statistical significance analysis, can be performed with uptake at a time t, uptake area up to a time t, the observed exchange rate constant or protection factors.

### Back-exchange correction

Despite efforts to minimize back-exchange in HDX-MS experiments^24–26^, there is always some percentage of the on-exchanged deuterium that exchanges back to hydrogen. For some types of analysis, such as the calculation of protection factors^27^, this needs to be corrected for. Prior to curve fitting, the user can choose whether to perform back-exchange correction^28^ or not. `HDfleX` allows for two methods of back-exchange correction, from: (1) an experimentally determined maximum deuteration (maxD) value, where the experimental maxD is included in the cluster data input file, and (2) a maximum deuteration value derived from the plateau value of the fitted uptake curve. The plateau maxD option can only be used on proteins and peptides that can undergo full exchange within the experimental time window.

### pH adjustment factor

The rate of HDX is affected by several factors, including pH and ionic strength^27,29,30^. When normalizing between different pH, `HDfleX` alters the experimental labelling timepoints according a pH adjustment factor, pH_adjust_factor_, according to Equation 2^31^, where pH_ref_ is the reference state pH and pH_exp_ is the experimental pH of the state being compared. pH_adjust_factor_ is subsequently multiplied to all the experimental labelling timepoints for that particular state.

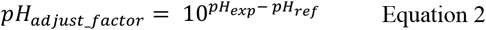

To test the pH adjustment, we used bradykinin in four different buffers (Table S1: Buffers 1-4). We chose bradykinin peptide (Arg1-Pro2-Pro3-Gly4-Phe5-Ser6-Pro7-Phe8-Arg9) as it has been found to adopt random-coil conformational states in aqueous media^32–34^. Due to its unstructured nature in aqueous medium, any changes in HDX in bradykinin between the conditions being compared can be attributed to the solution effects. HDX time courses on bradykinin with three replicates and seven timepoints were collected for each buffer condition being compared (see SI Methods for experimental details). The data was fitted and corrected for back-exchange as discussed above (Figure 2A). We first applied the pH adjustment to the four different conditions to normalize the solution effects among them (Figure 2B). If the solution effects differed only by pH, we would expect to see the uptake curves to perfectly overlay each other. However, our data showed that despite applying the pH correction, there were still differences up to 0.43 Da between the uptake curves, which is above the calculated global threshold of 0.09 Da for this peptide. This confirms that the salts used also influenced the intrinsic exchange kinetics^35^.

**Figure 2:**
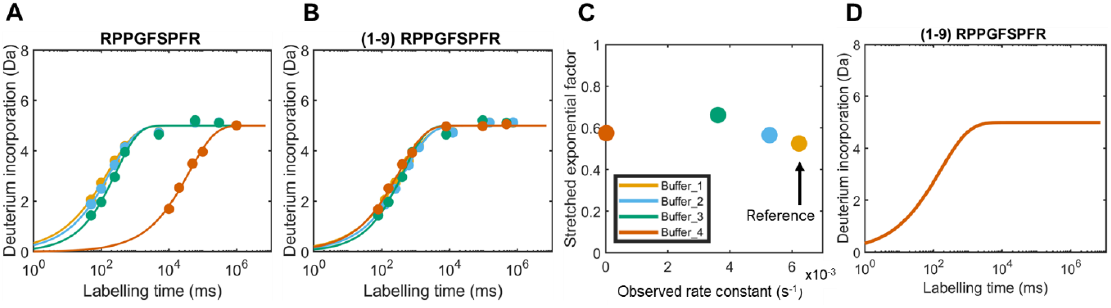
Calibrating the chemical exchange rate using bradykinin. (A) Bradykinin uptake curves in four states, back-exchange corrected and fitted in one exponential phase according to equation 1; (B) pH adjustment applied to the timepoints of bradykinin in each state prior to fitting; (C) Stretching factor *β* against the observed rate constant *k_obs_* for each buffer condition. (D) Empirical adjustment factor applied to bradykinin conditions, showing all four curves now perfectly overlaid (Δuptake = 0 Da).

### Empirical adjustment factor

As we saw in bradykinin, both pH and ionic composition can affect the intrinsic exchange rate^36^. In this case, it becomes difficult to discern the structural changes from the intrinsic exchange rate changes. To allow accurate comparison between conditions of different pH and ionic composition, we created a new empirical method which uses the unstructured peptide bradykinin to calibrate the intrinsic exchange rate effect.

Following back-exchange correction and curve-fitting, each state has two fitted parameters, *k_obs_* and *β*, associated with it. In Figure 2C, a plot of the stretching factor *β* against the observed rate constant *k_obs_* shows the *k_obs_* and *β* for each buffer condition. The first step is to choose a reference state (Buffer 1). A 2D scaling factor is determined for each coordinate in Figure 2C to transform to the reference coordinate. This scaling factor, consisting of a value for *k_obs_* and one for *β* is multiplied to the fitted parameters for all the buffer conditions. For bradykinin, this results in the uptake curves to perfectly overlay each other (Figure 2D), with differences of 0 Da between the curves, i.e., Δuptake = 0 Da. The empirical factor correction for each condition can then be applied to the uptake curves of the protein of interest, thereby normalizing the intrinsic exchange rate effects, allowing the structural effects to be clearly distinguished.

### Statistical significance testing

It is important to carry out statistical significance analyses in differential HDX experiments to determine significant differences across the protein between different experimental conditions. `HDfleX` takes the hybrid significance testing approach brought forward by Hageman and Weis for statistical significance testing^37^.

### Obtaining data distribution by separating the replicates

In the first stages of `HDfleX`, all the replicates were pooled together to perform the calculations. To calculate the global significance threshold for the HDX parameters (uptake area, observed rate constant and protection factors) except for the uptake, we need to know their distribution. Thus, it is necessary to perform curve fitting of different combinations of replicates. Here, we describe three methods by which `HDfleX` creates the distributions from the raw data for each peptide/fragment. SI Figure 5 shows a graphical explanation of each method. (1) Simple per replicate number combinations: At each timepoint, one replicate is chosen in numerical order until all the replicates have been processed. In cases of missing replicates at any particular timepoints, the latter will not include that replicate number in the curve fitting. The mean and SD of each of these distributions is calculated, (2) Random replicate combinations: At each timepoint, a random replicate is chosen from the set of replicates available. This is repeated *n* times, where *n* is the original number of replicates in the experiment. The mean and SD of each of these distributions is calculated, (3) Bootstrapping method of combinations: All the possible combinations of timepoints and replicates are initially calculated. A number *s* of random combinations is sampled uniformly from all the possible combinations, with replacement, using the MATLAB function `datasample`. Thereafter, 10,000 bootstrap data samples are drawn from *s* using the `bootstrp` function, and the mean and SD of each of these distributions is calculated.

### Global significance threshold

A global threshold is necessary to eliminate negligible differences that appear significant from the Welch’s t-test described in the next section, which would otherwise lead to false positives. Three methods of calculating the global significance threshold are available in `HDfleX`. The first one is based on the calculation of the confidence interval from experimental standard deviations (SD) as described previously by Hageman and Weis^37^. We have extended their method to include ETD fragments, to remove outliers and to be also used on the area under the curve, in addition to the uptake. The second method of calculating the global threshold, applying only to (1) and (2) data distribution methods above, initially involves calculating the pairwise differences between replicates within each state for the experimental data (SI Figure 6A). The pairwise difference is calculated rather than the difference between each replicate and the mean because we did not want any outliers in the dataset to skew the mean, which should be zero as the differences being calculated are within states. The outliers in the pairwise differences are optionally removed using the MATLAB function `isoutlier`^38^, with a method chosen by the user (mean, median, quartiles, grubbs, gesd), prior to fitting the pairwise replicate differences to a normal distribution. The upper confidence interval limit of this normal distribution is the global significance threshold. For the uptake area, the replicate differences and standard deviations are calculated from the integral of the fitted curves, whereas for the uptake, they are calculated from the experimental data points. When the bootstrapping method of combinations is selected (SI Figure 6B), the global threshold is calculated as simply being the upper limit of confidence interval of the all the means across the peptides and ETD fragments.

### Welch’s t-test

A Welch’s t-test is performed between the peptides and fragments of the two states being compared to give a p-value. The MATLAB function `ttest2` is used with the Satterthwaite’s approximation for the effective degrees of freedom^39^. To be significant, any differences between the peptides and fragments of the two states being compared must be above the global significance threshold and have a p-value lower than the alpha confidence level (e.g. p < 0.05)^37^.

### Data flattening over amino acid sequence

Instead of analyzing the HDX-MS on a per peptide/fragment basis, `HDfleX` flattens the data across all the available peptides or fragments to give high structural resolution HDX-MS data. The uptake area or uptake at time t at any peptide/fragment is divided by the number of exchangeable hydrogens (q). This number is subsequently applied across the amino acid residues covering that peptide as shown in SI Figure 7, except for the N-terminal residue and any prolines, which are fixed at 0. Note, for the ETD fragments, we attempted an intuitive subtraction of the consecutive fragments in the c/z series to obtain the deuterium uptake/area of highly resolved overlapping fragments, however this introduced large errors and was ultimately rejected (SI Figure 8).

### Visualizations of differences generated by HDfleX

`HDfleX` provides several ways to visualize the difference between states: difference and volcano plots at peptide resolution (un-flattened data) and amino acid residue resolution (flattened data). The differences between the means of the uptake area or uptake at any time t, are calculated by subtraction of one state from the other at each amino acid/peptide. Those differences are then plotted against the peptide/amino acid number to give a difference plot. Volcano plots for both flattened and un-flattened data are also generated for the p-value vs difference in uptake/uptake area^37^.

### 3D structural visualization of HDX differences

The differences calculated by `HDfleX` for uptake, uptake area, protection factor and observed rate constant are commonly visualized on the protein structure. `HDfleX` generates .pml script files at user-selected timepoints that can be run in PYMOL^40^.

### Calculation of intrinsic rates and protection factors

The intrinsic rates and protection factors were fitted and calculated as described previously^41^. Briefly, intrinsic uptake curves are generated, fitted into one- or two stretched exponentials. The intrinsic rate constant is extracted from the fitted parameters and the protection factor obtained using Equation 3, where *P_f_* is the protection factor, *k_int_* is the intrinsic amide hydrogen exchange rate constant from published values and *k_exp_* is the experimental rate constant from the experimental fits.

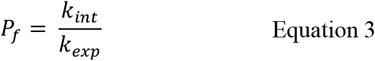

## RESULTS

To test `HDfleX`, we used datasets containing HDX-MS peptide and ETD information on the purified wild-type intrinsically disordered protein alpha-synuclein (aSyn) in various solution conditions (Table S1: Buffer conditions 1-4). Briefly, the millisecond timescale HDX was performed using a fully automated, millisecond HDX labelling and online quench-flow instrument, ms2min (Applied Photophysics, UK), connected to an HDX manager (Waters, USA). Mass spectrometry (MS) data were acquired on a Waters Synapt G2-Si Q-IM-TOF instrument with ETD capabilities. Both bottom-up and middle-down data were collected.

### Improved structural resolution when ETD fragments included with peptides

The results of `HDfleX` processing of HDX-MS data for aSyn shows an increase in structural resolution of MS data when ETD data are included with peptide data (Figure 3). Bottom-up data were combined with middle-down ETD data, resulting in a 50% increase (from 20 to 30) of single amino acids with fully resolved HDX data and a 150% and 230% increase in HDX data resolved at the dipeptides and tripeptides level, respectively (Figure 4). Overall, the merging of the ETD fragments and peptides have greatly improved the structural resolution. It was not possible to test any top-down or ECD data for this study, but these data should in principle be compatible with the `HDfleX` analysis to provide even higher structural resolution.

**Figure 3:**
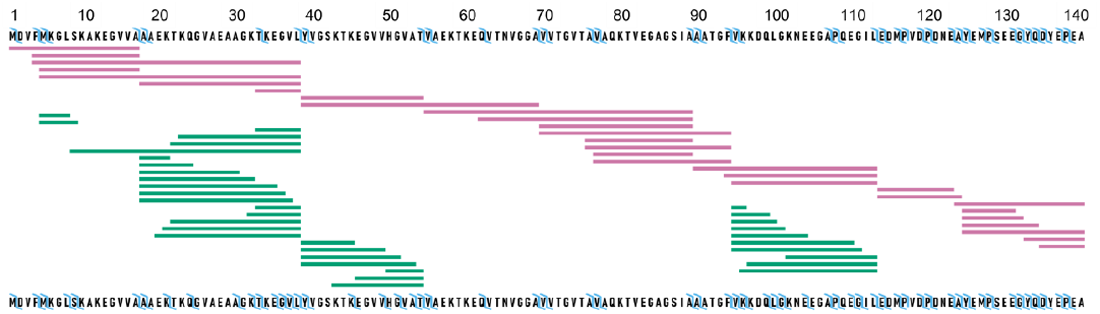
Inclusion of ETD fragments to peptide data improves structural resolution for sample data of aSyn HDX. Top sequence corresponds to peptide-only data (pink) and bottom sequence corresponds to peptide + ETD fragments data (green). The blue ticks represent the merged resolution obtained by combining peptides and c/z fragments.

**Figure 4:**
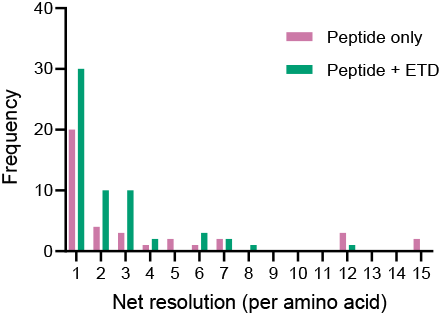
Comparison of the merged resolution between peptide only (pink) and peptide + ETD (green) data.

### Validation of global significance threshold methods

The importance of a global threshold for significance has recently been stressed by various groups for the correct statistical analysis of HDX-MS data^30,37,42^. `HDfleX` calculates the global threshold for the uptake (in Da) in the same way devised by Hageman and Weis^37^. For the uptake area global threshold, three methods are available: calculation of a confidence interval from (1) the standard deviations^37,43^, (2) the pairwise differences between replicates within states and (3) bootstrapped replicates.

To validate the three methods, we carried out a null experiment on the aSyn protein in Buffer 2 for four replicates and created six groups of two replicates from the same dataset (SI Figure 9). The difference in uptake area between the groups were calculated for all peptides, shown as scatter density plots in Figure 5. By definition, for a null experiment, the mean of the differences should be zero and this is what we observe. The spread of the differences is affected by outliers, as can be seen from the Null 3 dataset, for example. The presence of outliers alters the calculated global threshold for significant differences. `HDfleX` allows for the optional removal of outliers from the calculation of the global threshold using the MATLAB function `isoutlier`^38^. Without removing any outliers, we can see that for the 95% confidence intervals (Figure 5A), the bootstrap replicates and standard deviation-based confidence interval calculations result in no false positives. The other methods show significant differences due to false positives. Increasing the confidence levels to 99% (Figure 5B) removes false positives from the pairwise difference-based calculation using the simple per replicate data distribution method. However, the increase in confidence interval disproportionately alters the standard deviation and bootstrapping-based calculations and could result in false negatives, causing significant differences to be missed. Therefore, we consider that the bootstrap replicates and standard deviation-based calculations are highly robust at removing false positives whilst minimizing the risk of false negatives.

**Figure 5:**
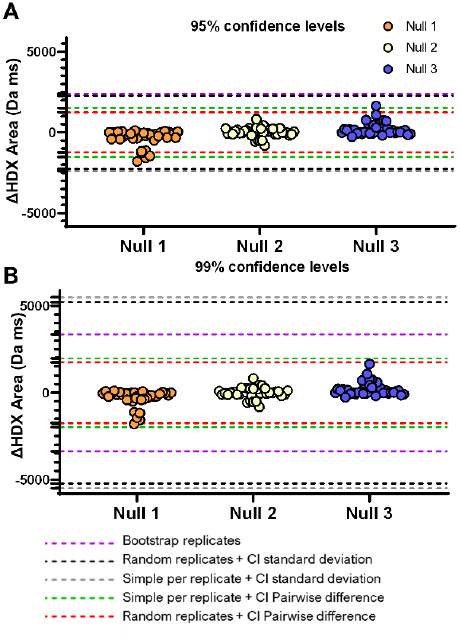
Comparison of the three methods of global threshold calculation on three null datasets derived from the protein aSyn in one condition only. The circles represent the difference in uptake between the two groups in the null dataset. Each null comparison was unique. The dashed lines represent the global thresholds calculated from each of the methods available in `HDfleX`. CI: confidence interval.

### Empirical adjustment as a novel normalization method for HDX-MS experiments

In order to normalize data between different buffer conditions (i.e. salts) that would alter intrinsic rates in a manner with no established solution, we used the unstructured peptide bradykinin^32,33^ as an internal standard. In this way, we aim to remove significant differences that are a result of buffer-derived changes to intrinsic rates and better differentiate those derived from changes to protein structural dynamics. The protein aSyn was on-labelled in the same buffers as bradykinin in separate experiments and the data kinetically fitted in `HDfleX` to obtain a rate constant (*k*) and stretch term (*β*), shown in Table S2. The empirical adjustment factors for *k* and *β* to convert to a reference state (here Buffer 1) are globally set values derived from bradykinin that are multiplied to the fitted parameters of each peptide and fragment of aSyn (SI Figure 10). It is important to note that this method is entirely empirical and has only been tested on the unstructured peptide bradykinin and intrinsically disordered protein aSyn in single-exponential phases, and therefore, we cannot assure it will work with more structured proteins.

### Intrinsic rate calculation and protection factors

`HDfleX` also calculates the theoretical intrinsic rates for peptides^41^ at the experimental pH and temperature. aSyn is incompatible with the Bai and Englander model for the calculation of intrinsic rates^27,44^ as all the peptides exchanged faster than the predicted rates. To demonstrate the ability of `HDfleX` to calculate intrinsic rates and protection factors, we analyzed previously described datasets on glycogen phosphorylase a and b (State_A and State_B, respectively)^41^. The uptake plot in Figure 6A shows the expected theoretical (black) and experimentally measured (blue and green) deuterium uptake for a selected peptide for both protein states, from which the *P_f_* is calculated. There is an upper limit of detection for *P_f_*, as determined by the measurable labelling times (for this dataset ln(*P_f_*) < 20). Therefore, a *P_f_* threshold can be set to discard unquantifiable, highly protected, slow-exchanging peptides. Following this filtering, 224 peptides were selected for the differential HDX analysis. To determine the significant differences between the two states, we performed hybrid significance testing as described earlier, the results of which are represented as a volcano plot (Figure 6B). Significant peptide uptake differences were found in 53% of the peptides and fall in the red shaded region. Furthermore, differences between State_A and State_B can be visualized as difference plots per peptide (Figure 6C and Figure 6D) and per amino acid residue (Figure 6E and 6F), with significant differences denoted with a red asterisk. We compared the uptake area difference plot per peptide generated by `HDfleX` to the butterfly plot generated by DynamX (SI Figure 11). Multiple features were found to be (in)significant by `HDfleX`, which would have the opposite evaluation without fitting and statistical analysis. Here, we show an important distinction when using protection factors is that ‘hidden’ features with low uptake but with high protection can be detected (shown as blue shaded region in Figure 6C and 6D).

**Figure 6:**
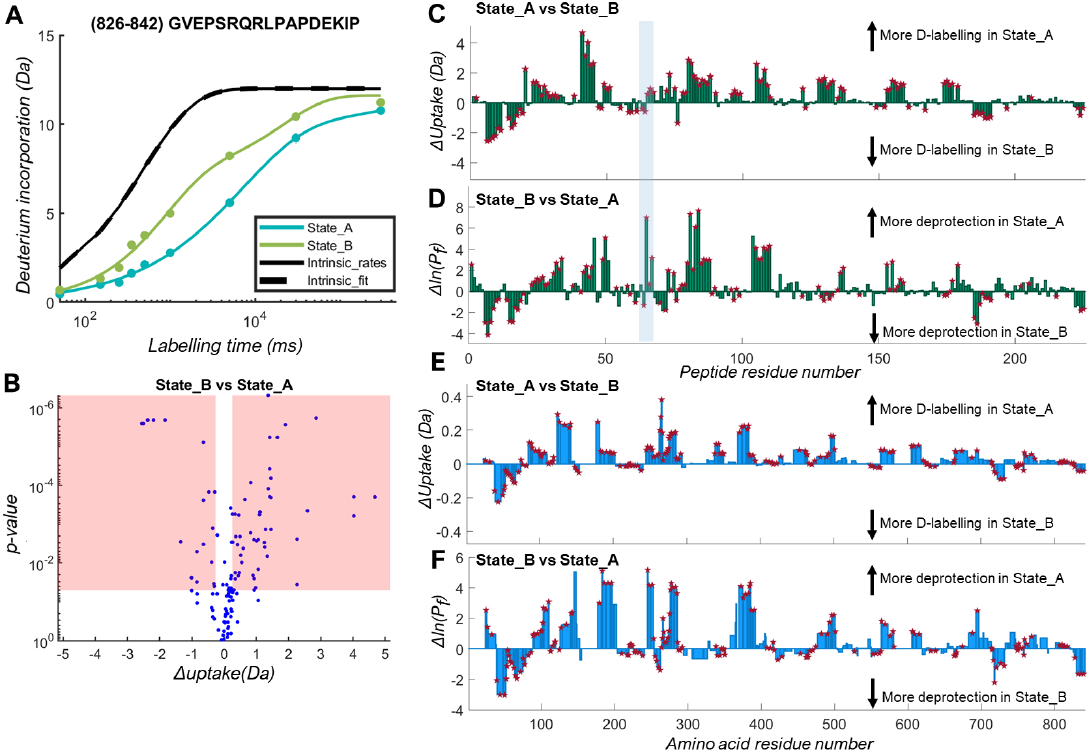
Glycogen phosphorylase HDX-MS data analyzed with `HDfleX`. (A) Example deuterium uptake plot showing the experimental and theoretical fitting of the glycogen phosphorylase State_A and State_B; (B) Volcano plot representation of the hybrid significance testing comparison of State_A vs State_B; (C) Uptake difference plot of State_A vs State_B at peptide resolution; (D) ln(*P_f_*) difference plot of State_A vs State_B at peptide resolution; (E) Uptake difference plot of State_A vs State_B at amino acid residue resolution; (F) ln(*P_f_*) difference plot of State_A vs State_B at amino acid residue resolution. Significant differences are denoted by a red asterisk.

## CONCLUSION

In this paper, we presented `HDfleX`, a new standalone application for the analysis of HDX-MS data postprocessing, allowing the combination of data at different levels: peptides and ETD fragments. Other currently available programs do not allow the combination of data at different fragmentation levels, limiting the analysis of HDX-MS data and the resulting structural resolution. With `HDfleX`, we saw an improvement in structural resolution of the HDX-MS data for areas where ETD data was collected, up to a 50% increase in single residues.

We have implemented a robust significance testing method, adapting the hybrid significance testing approach by Hageman and Weis^37^. The adjustments we included, with respect to outlier removal make the calculation of the global significance testing more robust to the presence of outliers. Throughout, we use the ‘uptake area’, as opposed to the more commonly used ‘uptake’, as our main parameter. Other groups have proposed the use of uptake area in differential HDX studies^37,45^, but with no global threshold calculation method available due to the added complexity of the different combinations of replicate data points – a problem now solved by `HDfleX`. As such, not only did we expand the hybrid significance testing method to work on uptake areas, but we created two new methods of calculating the global threshold for the uptake area (based on replicate variability and bootstrapping). This enables users the option to use uptake area easily and effortlessly in their analyses, whilst maintaining robust elimination of false positives and false negatives.

`HDfleX` generates the theoretical intrinsic uptake plots and protection factor, based on the intrinsic rates devised by Bai et al^27^. We also introduced a novel empirical approach to normalize between various conditions (e.g., salts) and for proteins (e.g., intrinsically disordered proteins) in a way not accounted for by established correction^27^. We tested this approach on the unstructured peptide standard bradykinin and the intrinsically disordered aSyn. We hope that this new method, when used with scrutiny, will expand the types of differential HDX-MS experiments that can be done, as at present, it is rare to see comparisons between factors that affect the intrinsic rate (such as pH, temperature, ionic strength)^31,46^.

`HDfleX` is available to download from http://hdl.handle.net/10871/127982 and does not require a MATLAB license to use.

## ASSOCIATED CONTENT

The Supporting Information is available free of charge on the ACS Publications website.

Text, Additional method details (Bradykinin HDX-MS experimental details); Figures, flowcharts describing import and fitting stage, data flattening and statistical significance analysis; plateau back-exchange determination; methods of data distribution by separating the replicates; error distribution of the entire dataset; schematic of data flattening process; testing weights for data flattening; deuterium uptake plots for null experiment on aSyn; application of empirical adjustment to an aSyn peptide; comparison of difference plots generated by DynamX vs `HDfleX`; Tables, Bradykinin buffer chemical compositions; determination of empirical adjustment factor from bradykinin fitted parameters.

## ACKNOWLEDGEMENTS

NS is funded by the University Council Diamond Jubilee Scholarship. JJP and MK are supported by a UKRI Future Leaders Fellowship [Grant number: MR/T02223X/1]. We are grateful for fruitful discussions with Dr S. Subramanian.

