## Supplementary_Information for "HDfleX: Software for flexible high structural resolution of hydrogen/deuterium-exchange mass spectrometry data"

### SI Materials

All reagents and chemicals were purchased from Sigma-Aldrich, USA, unless mentioned otherwise.

### SI Methods

*Bradykinin HDX-MS experimental details*

0.5 µL of 10 mg/mL bradykinin peptide in dimethyl sulfoxide was added to 500 µL of each of the equilibration buffers (Table S1: Buffers 1-4) and allowed to equilibrate at room temperature for 1 hour. HDX for timepoints under 300 s was performed using a fully automated, millisecond HDX labelling and online quench-flow instrument, ms2min (Applied Photophysics, UK). HDX for timepoints above 300 s were performed using a CTC PAL sample handling robot (LEAP Technologies, USA). The ms2min or CTC PAL robot was connected to an HDX manager (Waters, USA). Mass spectrometry (MS) data were acquired on a Waters Synapt G2-Si Q-IM-TOF instrument. Three technical replicates and seven timepoints were collected. For Buffers 1-3, the timepoints were: 50 ms, 100 ms, 250 ms, 500 ms, 5000 ms, 60 s, 300 s. For Buffer 4, the timepoints were: 10 s, 20 s, 50 s, 100 s, 998 s, 11972 s, 59858 s. Deuterium incorporation was determined in DynamX 3.0 (Waters, USA). Back-exchange correction was performed using the uptake plateau determination method in `HDfleX`.

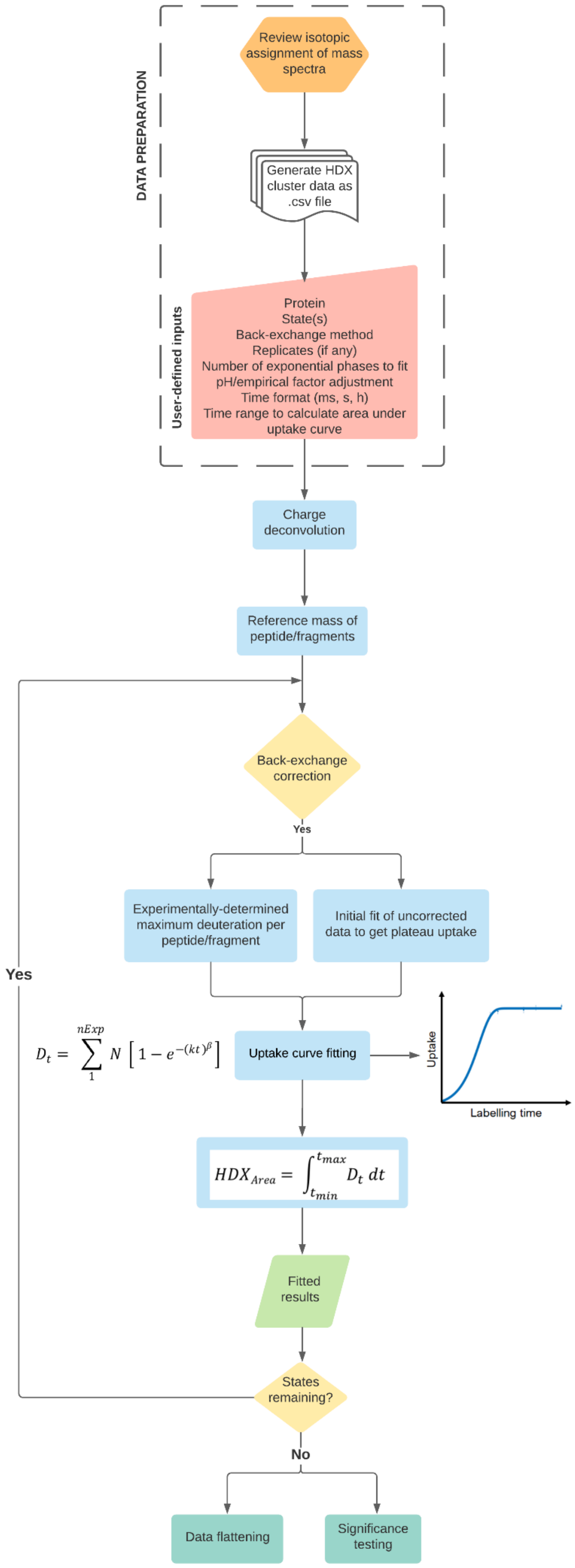


**SI Figure 1:** Flowchart describing the import and fitting stage of HDfleX

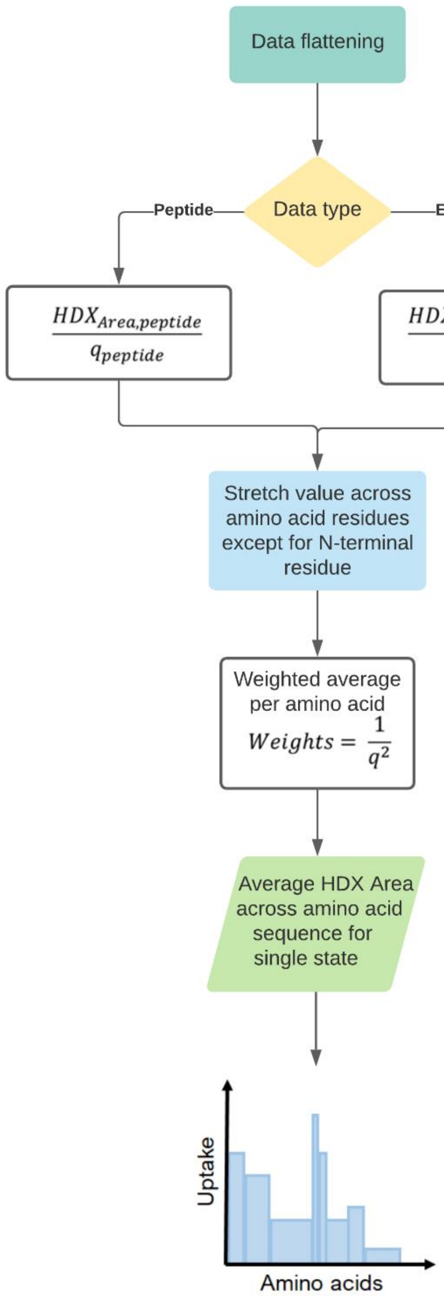


**SI Figure 2**: Flowchart describing the data flattening stage of HDfleX.

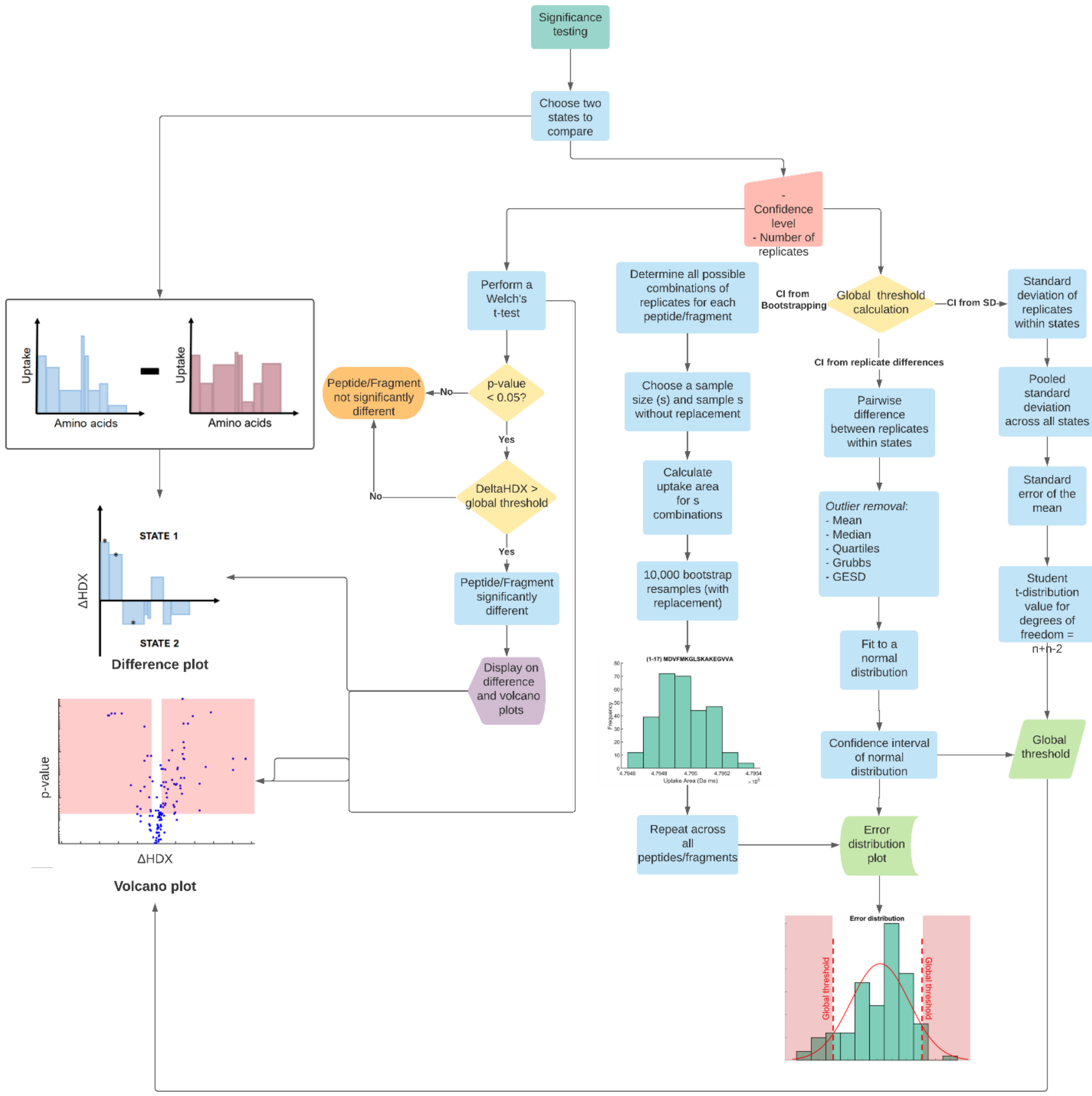


**SI Figure 3**: Flowchart describing the statistical significance analysis stage of HDfleX.

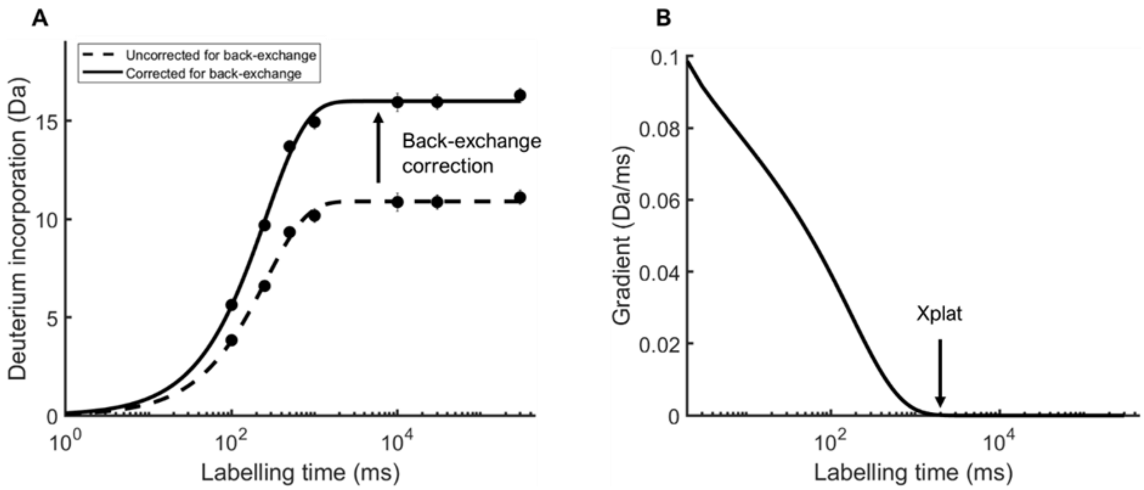


**SI Figure 4**: Plateau back-exchange determination. (A) Back-exchange correction shifts the uncorrected uptake curve (dotted) to the corrected curve (solid) for an example peptide; (B) The gradient of the fitted uncorrected curve is determined per time unit. The time at which the gradient is < 0.0001 is called Xplat and the uptake at Xplat is the maxD.

**Table S1: Buffer compositions.**

| Ion/Additive | Buffer 1 (mM) | Buffer 2 (mM) | Buffer 3 (mM) | Buffer 4 (mM) | Buffer 5 (mM) |
|---|---|---|---|---|---|
| **$Na^+$** | 0 | 143 | 15 | 20 | 0 |
| **$K^+$** | 0 | 4 | 140 | 60 | 0 |
| **$Ca^{2+}$** | 0 | 2.5 | 0.0001 | 0 | 0 |
| **$Mg^{2+}$** | 0 | 0.7 | 10 | 0 | 0 |
| **TCEP** | 0 | 0 | 0 | 0 | 1 |
| **pH** | 7.4 | 7.4 | 7.2 | 4.9 | 7.0 |
| **Buffer system** | 20 mM Tris | 20 mM Tris | 20 mM Tris | 20 mM citrate | 40 mM Tris |

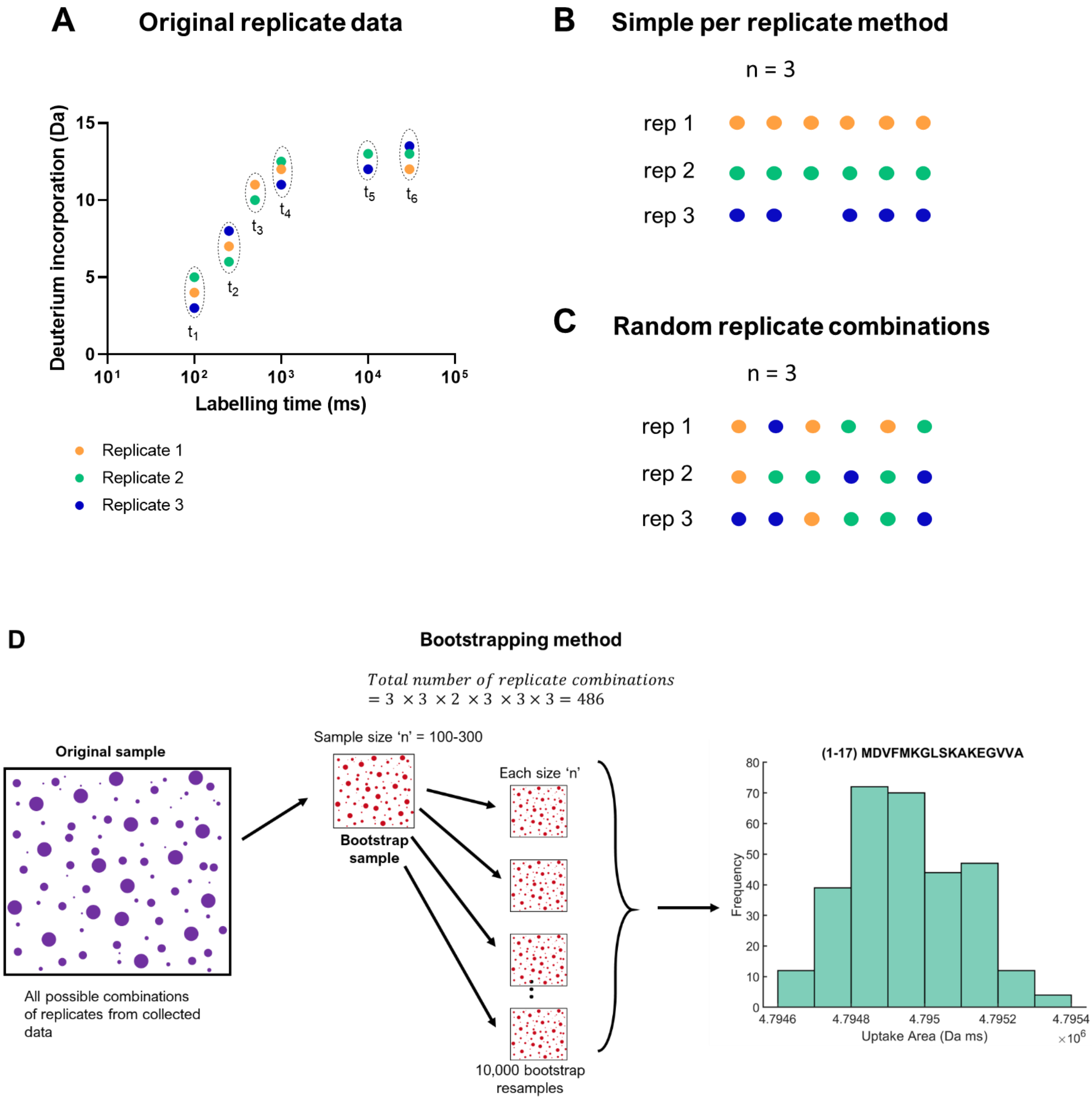


**SI Figure 5:** Methods of data distribution by separating the replicates; **(A)** Original replicate data for a chosen peptide with data points coloured as follows: replicate 1 = orange, replicate 2 = green, replicate 3 = blue. Each timepoint is encircled with broken lines; **(B)** Simple per replicate method: one replicate is chosen at each timepoint in numerical order until all experimental replicates have been processed; **(C)** Random replicate combinations method: A random replicate data point from the available replicates is chosen at each timepoint and this is repeated n times; **(D)** Bootstrapping method: All possible combinations of replicates at all timepoints are fitted and the uptake area is calculated. A bootstrap sample size of 300 was chosen here, and this sample size is resampled with replacement 10,000 times. The resulting uptake area distribution is shown for the chosen peptide, from which the mean and standard deviation can be determined.

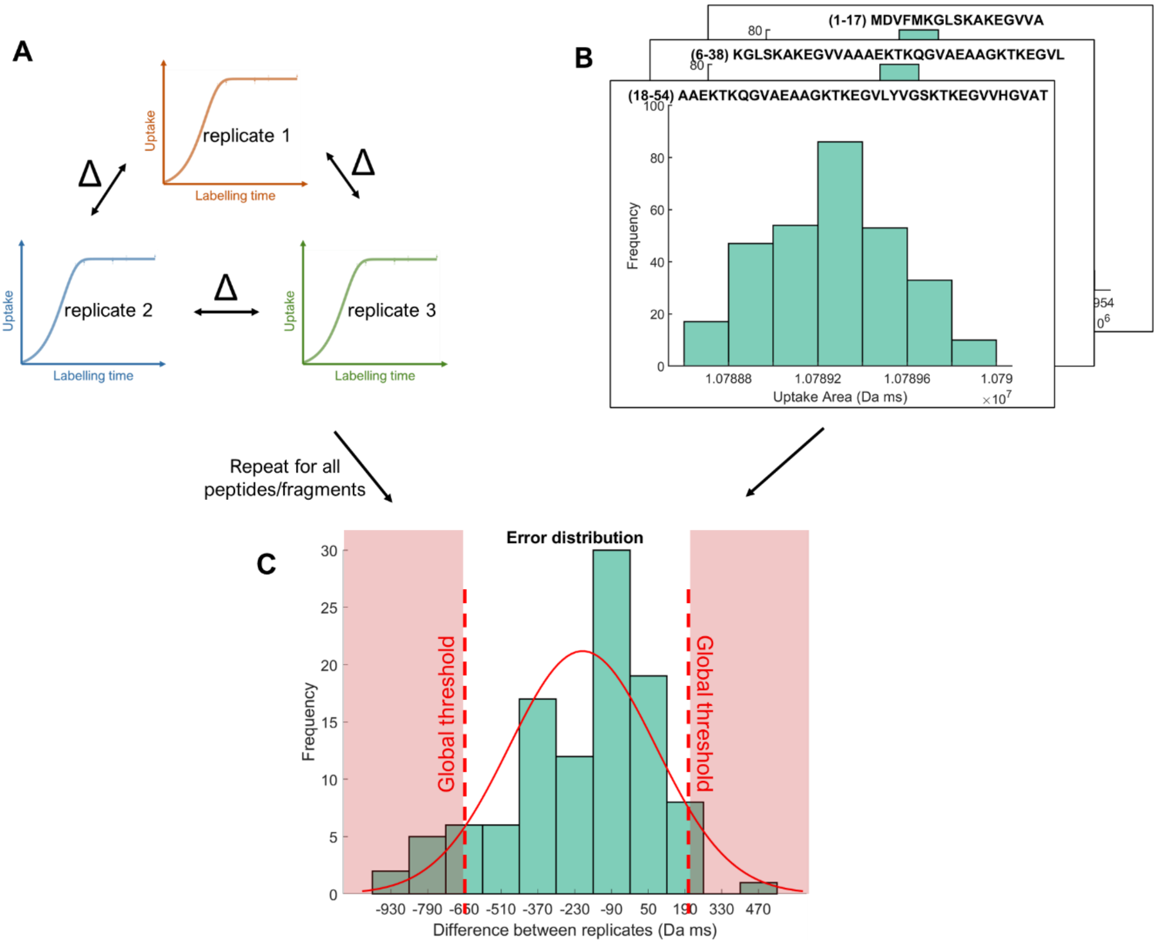


**SI Figure 6:** Error distribution of entire dataset using two different methods. (A) When the simple per replicate or random replicate combinations method of data distribution are used, the pairwise difference between the replicates for a peptide/fragment within each state is calculated. These pairwise differences are subsequently fitted with a normal distribution; (B) When using bootstrapping method of data distribution, the error distribution is determined from the maximum confidence interval of the mean across all peptides and fragments. (C) Resulting error distribution: the red shaded region represents differences that are large enough to be significant.

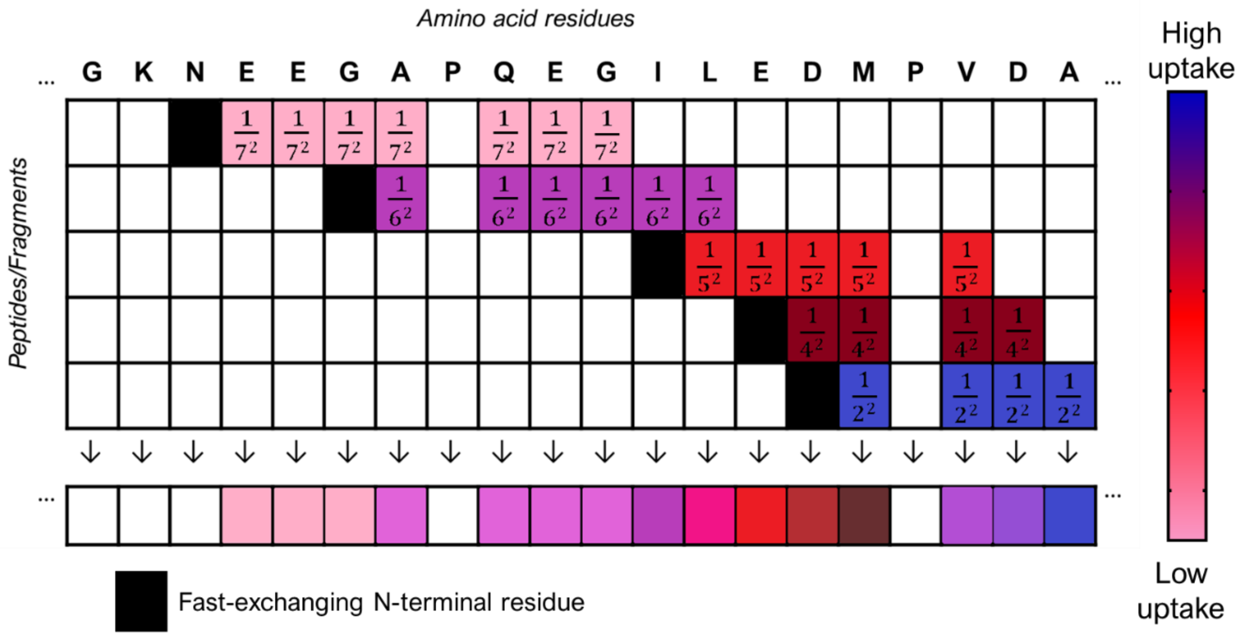


**SI Figure 7:** Schematic of the data flattening process adapted from Keppel & Weis, 2014[3]. The different colours represent different uptake/area values, with blanks representing no uptake/area, and the black squares representing rapidly exchanging N-terminal amide hydrogens. The bottom grid shows the flattened data which is obtained by calculating a weighted average of the uptake/area at each amino acid (per column) given by Equation 3.

**Testing various weights for the peptides and fragments, including subtraction of consecutive ETD fragments**

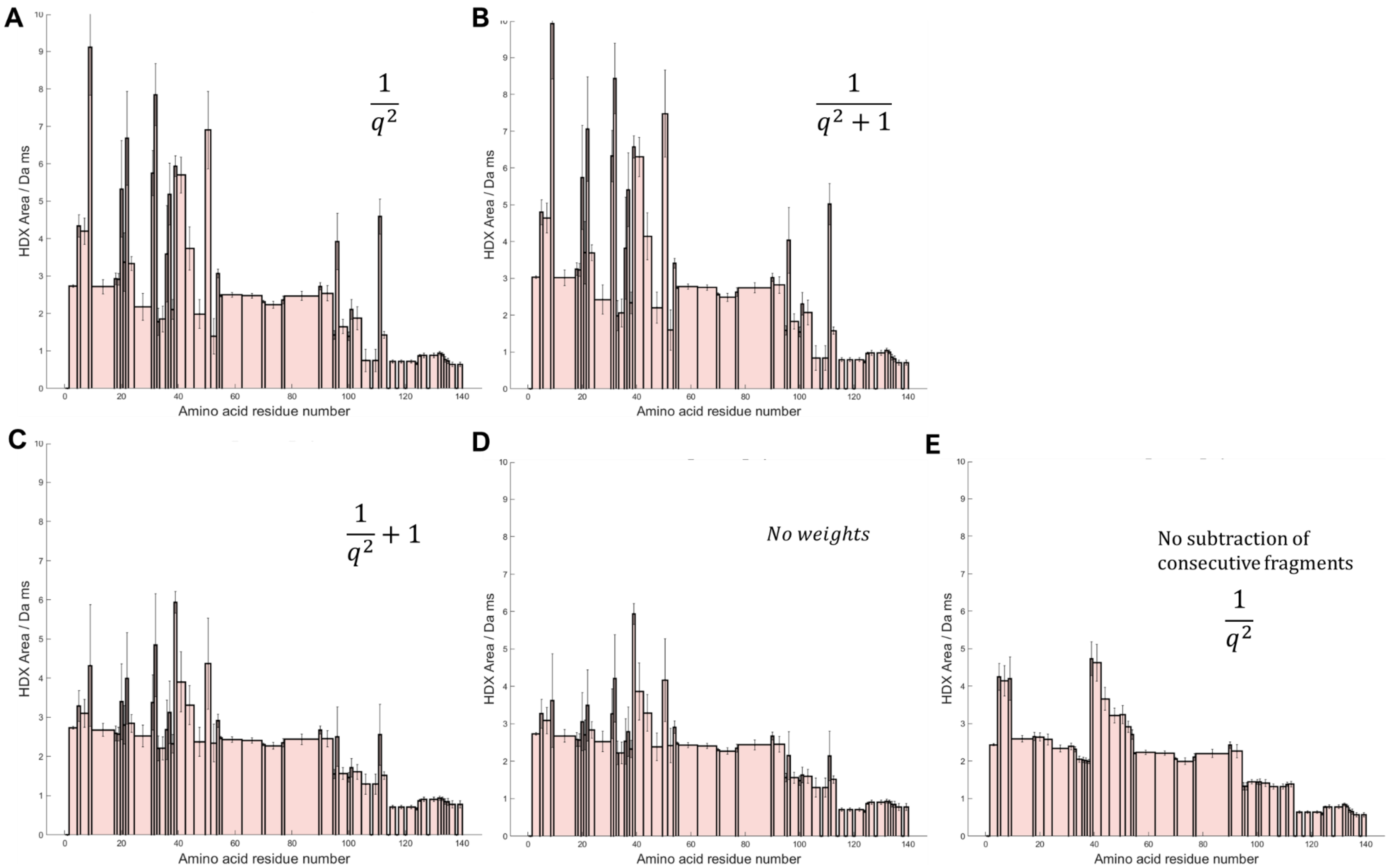


**SI Figure 8:** A range of weights were tested for the flattening of the peptides and ETD fragments; the weights are shown alongside the plots. In (A)-(C), the flattening was performed using the weights shown, as well as the subtraction of consecutive ETD fragments; (D) No weights were used but subtraction of consecutive ETD fragments was still performed; (E) Weight of 1/q^2 was used and no subtraction of the fragments performed.

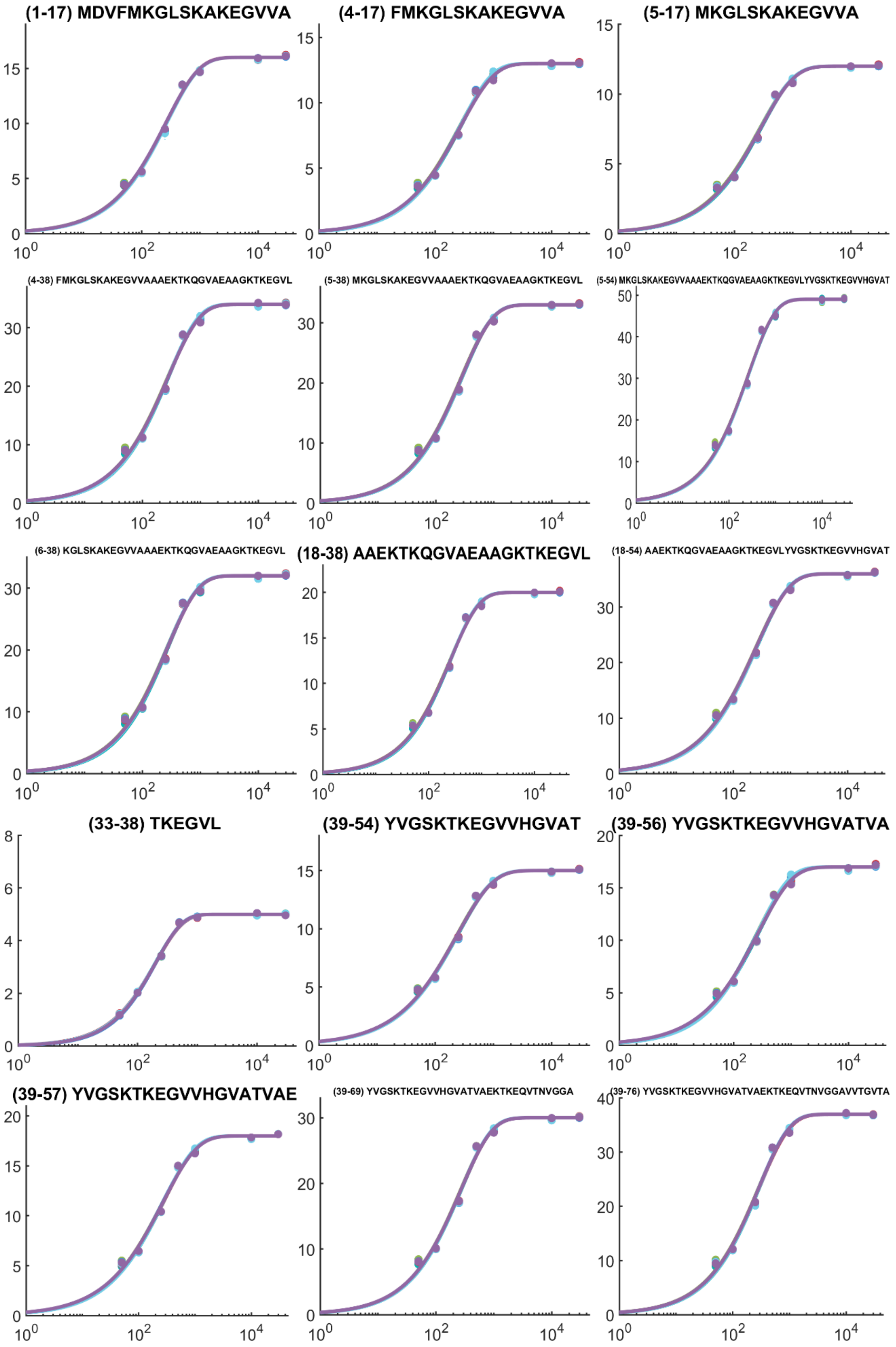
(1-17) MDVFMKGLSKAKEGVVA
(4-17) FMKGLSKAKEGVVA
(5-17) MKGLSKAKEGVVA
(4-38) FMKGLSKAKEGVVAAAEKTKQGVAEAAGKTKEGVL
(5-38) MKGLSKAKEGVVAAAEKTKQGVAEAAGKTKEGVL
(5-54) MKGLSKAKEGVVAAAEKTKQGVAEAAGKTKEGVLYVGSKTKEGVVHGVAT
(6-38) KGLSKAKEGVVAAAEKTKQGVAEAAGKTKEGVL
(18-38) AAEKTKQGVAEAAGKTKEGVL
(18-54) AAEKTKQGVAEAAGKTKEGVLYVGSKTKEGVVHGVAT
(33-38) TKEGVL
(39-54) YVGSKTKEGVVHGVAT
(39-56) YVGSKTKEGVVHGVATVA
(39-57) YVGSKTKEGVVHGVATVAE
(39-69) YVGSKTKEGVVHGVATVAEKTKEQVTNVGGA
(39-76) YVGSKTKEGVVHGVATVAEKTKEQVTNVGGAVVTGVTA

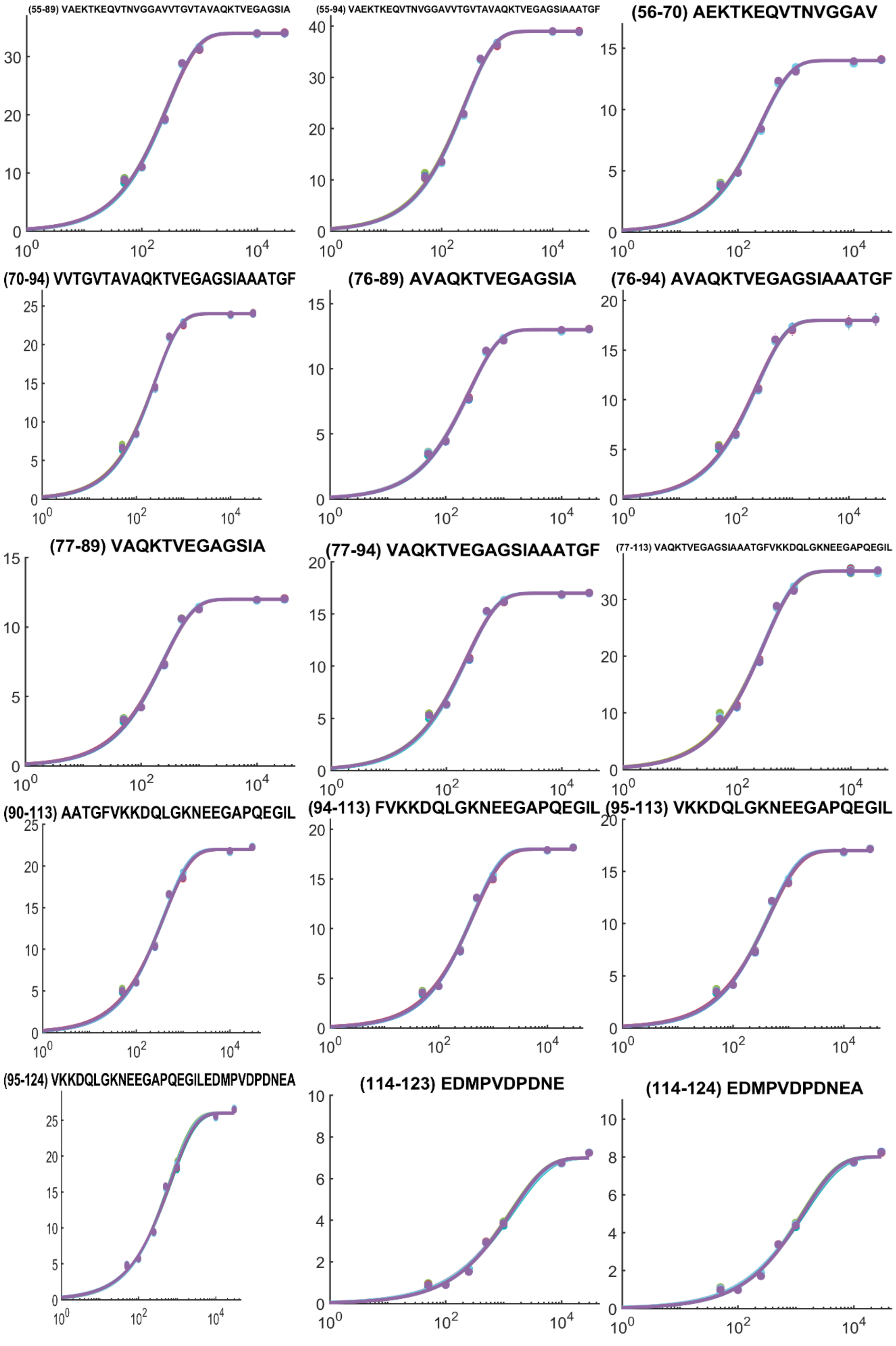
(55-89) VAEKTKEQVTNVGGAVVTGVTAVAQKTVEGAGSIA
(55-94) VAEKTKEQVTNVGGAVVTGVTAVAQKTVEGAGSIAAATGF
(56-70) AEKTKEQVTNVGGAV
(70-94) VVTGVTAVAQKTVEGAGSIAAATGF
(76-89) AVAQKTVEGAGSIA
(76-94) AVAQKTVEGAGSIAAATGF
(77-89) VAQKTVEGAGSIA
(77-94) VAQKTVEGAGSIAAATGF
(77-113) VAQKTVEGAGSIAAATGFVKKDQLGKNEEGAPQEGIL
(90-113) AATGFVKKDQLGKNEEGAPQEGIL
(94-113) FVKKDQLGKNEEGAPQEGIL
(95-113) VKKDQLGKNEEGAPQEGIL
(95-124) VKKDQLGKNEEGAPQEGILEDMPVDPDNEA
(114-123) EDMPVDPDNE
(114-124) EDMPVDPDNEA

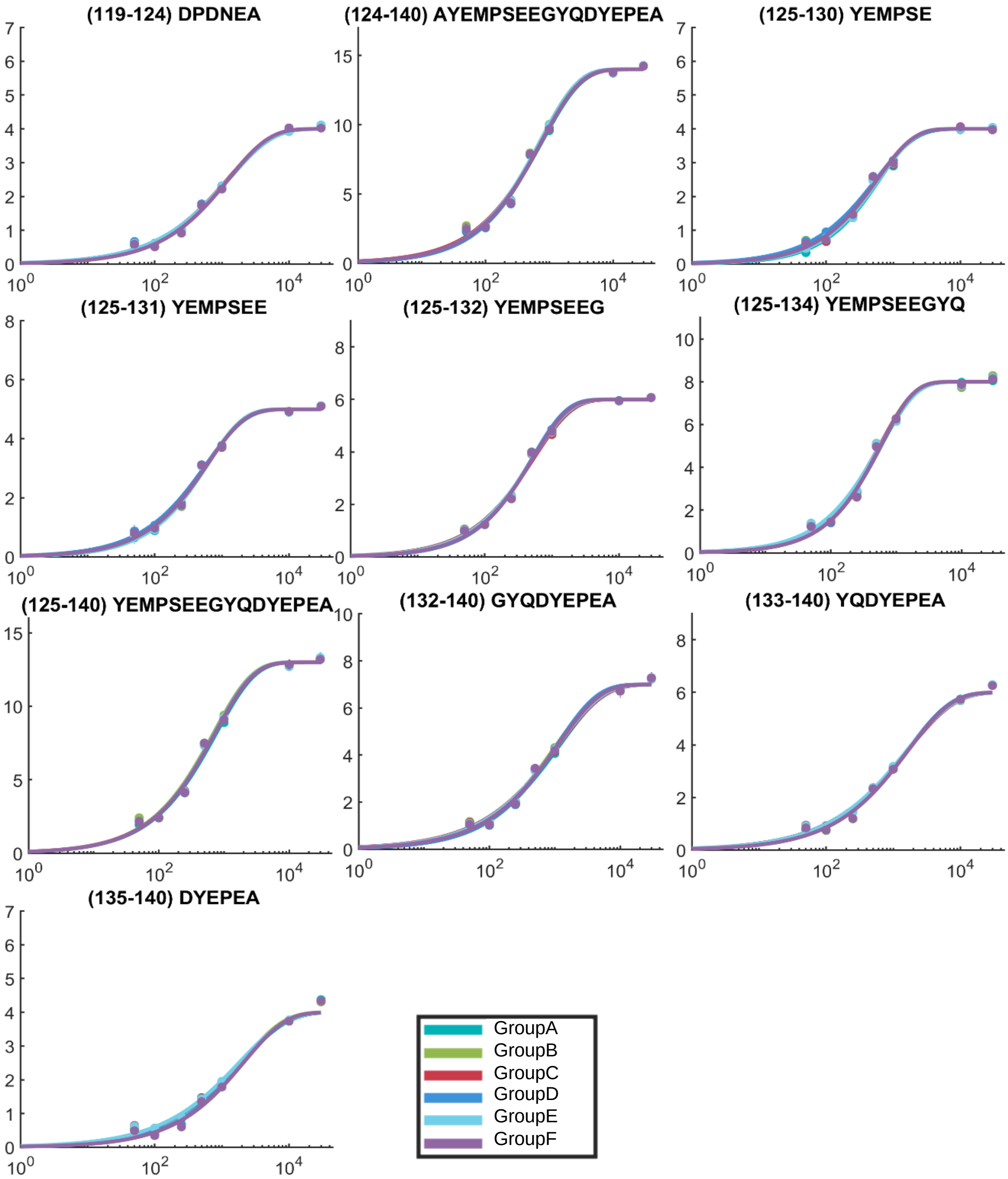


**SI Figure 9**: Deuterium uptake plots for the aSyn null experiment.

**Table S2**: Fitted parameters for the uncorrected and corrected states of bradykinin, and the empirical adjustment factor applied to each fitted parameter.

| States | Uncorrected fitting parameters | | Corrected fitting parameters | | Empirical adjustment factor | |
|---|---|---|---|---|---|---|
| | *k* | *β* | *k* | *β* | *k* | *β* |
| Buffer 1 | 0.0062 | 0.5247 | 0.0062 | 0.5247 | 1 | 1 |
| Buffer 2 | 0.0053 | 0.5651 | 0.0062 | 0.5247 | 1.179 | 0.9284 |
| Buffer 3 | 0.0036 | 0.6616 | 0.0062 | 0.5247 | 1.722 | 0.7931 |
| Buffer 4 | $2.49 \times 10^{-5}$ | 0.5749 | 0.0062 | 0.5247 | 250.1 | 0.9126 |

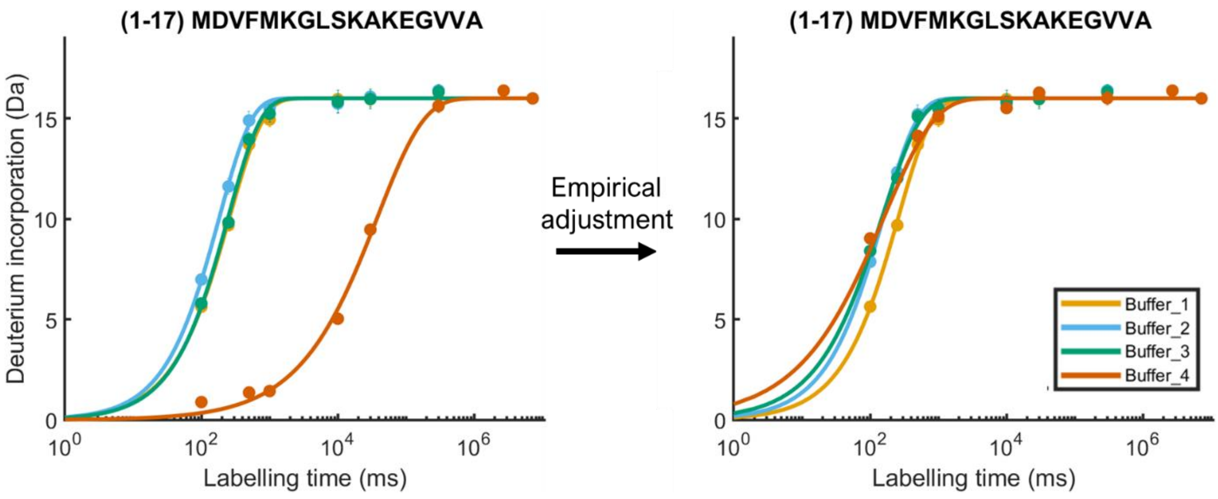


**SI Figure 10:** Application of empirical adjustment on aSyn peptide 1-17 for buffers 1-4, with Buffer_1 chosen as the reference condition.

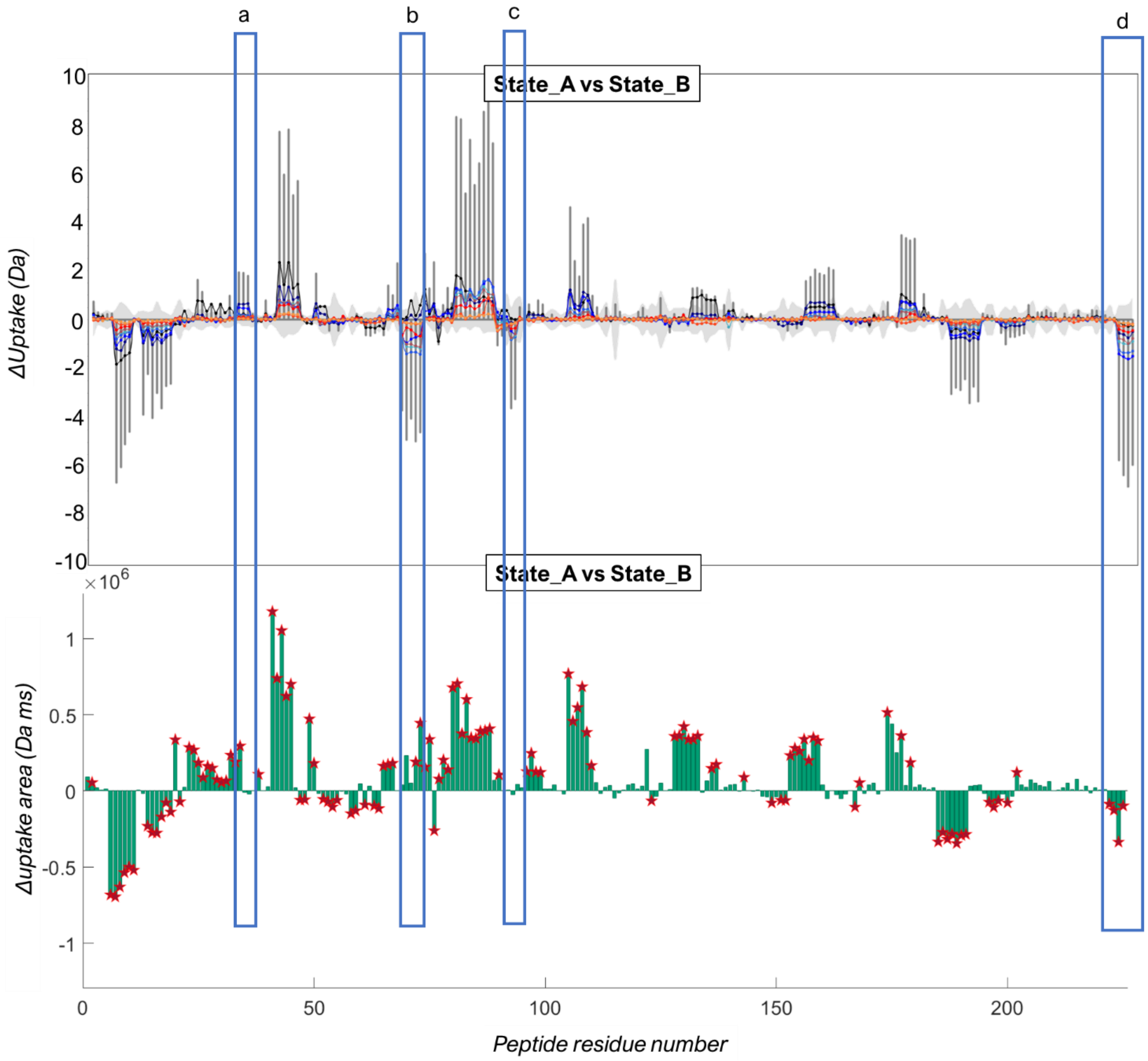


**SI Figure 11**: DynamX and `HDfleX` result in distinct difference plots per peptide residue for the comparison of glycogen phosphorylase State_A and State_B. *Top*: the difference 'butterfly' plot output by DynamX shows the difference in the sum of deuterium incorporated over all timepoints up to 300 s between the two states and includes no hybrid significance analysis. *Bottom*: the difference plot output by `HDfleX` shows the difference in uptake area up to 300 s and includes hybrid significance analysis. Drastically different regions between the two comparisons are encased in a blue box, and labelled a-d.
